# Sodium nitrophenolates as a potential tool for the management of grapevine trunk diseases

**DOI:** 10.64898/2026.10.21.760630

**Authors:** Jana Čechová, Kateřina Štůsková, Pavla Přinosilová, Linda Holcová, Dana Homolová, Milan Špetík, Petr Babuška, Ondřej Dvořák, Aleš Eichmeier

## Abstract

Fungal trunk diseases represent a major constraint on the production of grapevine and other perennial crops, while effective curative control methods remain limited. This study evaluated the antifungal and defence-modulating effects of the sodium nitrophenolate based product Atonik® against selected fungi associated with trunk diseases. *In vitro* efficacy was tested against ten fungal pathogens, and its effect against *Eutypa lata* was further evaluated in grapevine under *in planta* conditions. Atonik® inhibited mycelial growth in a pathogen and concentration dependent manner, with the strongest effects at the highest tested concentration observed against *Cadophora luteo-olivacea*, *E. lata*, and *Phaeoacremonium minimum*. In grapevine, *E. lata* caused a mean lesion length of 48.38 mm, whereas Atonik® treatment reduced it to 12.29 mm. Absolute qPCR indicated higher *VvSOD* transcript abundance in treated plants. RNA-seq analysis identified 168 genes associated with infection and 43 genes associated with treatment. The results suggest that sodium nitrophenolates may reduce wood necrosis caused by *E. lata* through a combination of direct antifungal activity and modulation of grapevine defence responses.

## Introduction

Fungal trunk diseases represent a complex group of destructive woody plant diseases affecting perennial crops and ornamental species worldwide ^1^. In grapevine, these diseases are collectively referred to as grapevine trunk diseases (GTDs) and are considered among the most serious threats to the sustainability and productivity of modern viticulture ^2^. Similar disease complexes also occur in fruit trees and ornamental nursery plants ^3^, where they contribute to wood necrosis, vascular dysfunction, dieback, decline, reduced yield, and, in severe cases, plant death. The economic impact of trunk diseases is particularly significant because symptoms often develop progressively over several years and effective curative treatments are currently unavailable ^2,4^.

GTDs are associated with a broad diversity of fungal pathogens colonizing woody tissues. More than 130 fungal species have been reported as causal or associated agents of GTDs, including representatives of the genera *Eutypa*, *Phaeoacremonium*, *Phaeomoniella*, *Cadophora*, *Dactylonectria*, *Diplodia*, *Neofusicoccum*, *Botryosphaeria*, *Diaporthe*, and *Fomitiporia*. These fungi may infect plants through pruning wounds, propagation material, root systems, or natural openings and can persist asymptomatically before disease expression ^5^. Among them, *Eutypa lata*, *Phaeoacremonium minimum*, *Phaeomoniella chlamydospora*, *Diplodia seriata*, and *Neofusicoccum* spp. are frequently reported as highly relevant grapevine trunk disease-associated pathogens due to their wide distribution, pathogenicity, and association with severe wood necrosis, canker formation, vine decline, and dieback symptoms ^2,4,6,7,8^.

Current management of trunk diseases relies mainly on preventive approaches, including the use of healthy planting material, pruning wound protection, sanitation measures, biological control agents, and cultural practices that reduce plant stress. However, the increasing pressure to reduce conventional pesticide inputs has stimulated interest in alternative compounds capable of limiting pathogen growth and/or enhancing plant defence responses ^2,9,10,11,12,13^. In this context, sodium nitrophenolates have gained attention as compounds with potential effects on plant physiology, stress tolerance, and defence-related metabolism ^14,15,16^. These compounds are commonly used in products containing sodium 5- nitroguaiacolate, sodium o-nitrophenolate, and sodium p-nitrophenolate, but their direct effects on fungal trunk disease-associated pathogens and their influence on host responses remain insufficiently understood.

Nitrophenols are phenolic compounds with the general formula HOC₆H₅₋ₓ(NO₂)ₓ, in which one or more nitro groups are attached to carbon atoms of the benzene ring. Accordingly, they are classified as mono-, di-, or tri-nitrophenols. Mononitrophenols occur as three structural isomers, which differ in the position of the carbon atom bearing the nitro group. These include ortho-nitrophenol, also referred to as 2-nitrophenol, in which the hydroxyl and nitro groups are located on adjacent carbon atoms; meta-nitrophenol, or 3-nitrophenol; and para-nitrophenol, or 4-nitrophenol ^17,18^. Nitrophenols exhibit acidic properties, and their conjugate bases are referred to as nitrophenolates. Although nitrophenolates have been successfully used in the cultivation of most agricultural crops for many years, their precise mode of action has not yet been fully elucidated ^19^. In general, they are reported to prolong the activity of auxins through inhibition of IAA oxidases and modulation of abscisic acid effects, enhance nutrient uptake by accelerating cytoplasmic streaming and assimilate uptake and to increase nitrate reductase activity. Przybysz et al.^14^ published a review summarizing the mode of action of nitrophenol-based biostimulants. They investigated the effects of Atonik® on oilseed rape, field cucumber, and the model plant Arabidopsis thaliana, which was deliberately grown under stress conditions. They concluded that nitrophenolates affect plants at all levels of their structure and function, from leaves and whole plants to individual organs and cells, as well as physiological and biochemical processes. In particular, it promotes generative growth and plant development and increases biomass production through stimulation of the photosynthetic apparatus ^14^.

The present study evaluates the *in vitro* inhibitory activity of sodium nitrophenolates against selected fungal trunk disease-associated pathogens, namely *Dactylonectria torresensis*, *Cadophora luteo-olivacea*, *Eutypa lata*, *Phaeoacremonium minimum*, *Phaeomoniella chlamydospora*, *Diaporthe ampelina*, *Fomitiporia mediterranea*, *Diplodia seriata*, *Botryosphaeria dothidea*, and *Neofabraea kienholzii*. In addition, the effect of these compounds on the virulence of *E. lata* was assessed under *in planta* conditions using grapevines. Particular attention was also given to the expression of *VvSOD*, a gene encoding superoxide dismutase, in order to determine whether sodium nitrophenolates may influence oxidative stress related defence responses in grapevine. This study therefore provides insight into both the direct antifungal potential of sodium nitrophenolates and their possible role in modulating grapevine responses to *Eutypa lata*.

## Results

### Inhibitory activity of Atonik® in vitro and in planta

The inhibition percentages are presented in Table 1. Atonik® has statistically significant effect on almost all tested trunk pathogens in the basic concentration (0.75% vol. 1:150) except of *F. mediterranea* isolate FOP1. Regarding the used spectra of used dilutions Atonik® showed the most persistent activity across dilutions against BD (2-1) and the highest percentace levels reached the efficiency against CLO (H20). The lowest efficiency was observed against FM (FOP1) and DT (CZ2).

**Table 1.** Statistically significant differences between each treatment and the control (fungus without Atonik® treatment), determined using Tukey’s test, are indicated by asterisks (*). DT – *Dactylonectria torresensis*, CLO – *Cadophora luteo-olivacea*, EL – *Eutypa lata*, PM – *Phaeoacremonium minimum*, PC – *Phaeomoniella chlamydospora*, DA – *Diaporthe ampelina*, FM (FOP1) – *Fomitiporia mediterranea*, DS – *Diplodia seriata*, BD – *Botryosphaeria dothidea*, NK – *Neofabraea kienholzii*

| Substance | Inhibition (%) |  |  |  |  |  |  |  |  |  |
| --- | --- | --- | --- | --- | --- | --- | --- | --- | --- | --- |
|  | Pathogen |  |  |  |  |  |  |  |  |  |
|  | DT (CZ2) | CLO (H20) | EL (CS 12 27 4 KO) | PM (ESP1) | PC (CS 61/14/2 ZN H55) | DA (H23) | FM (FOP1) | DS | BD (2-1) | NK |
| AT 0.75% vol. 1:150 | 33.13* | 74.62* | 75.62* | 70.92* | 58.45* | 50.30* | 25.62 | 23.03* | 34.05* | 51.15* |
| AT 10× 0.075% vol. 1:1500 | 0.00 | 32.32* | 14.51 | 30.50 | 31.51 | 25.70 | 0.00 | 5.37 | 25.76* | 11.90 |
| AT 100× 0.0075% vol. 1:15000 | 0.00 | 32.97* | 18.36 | 34.40 | 19.63 | 18.13 | 0.00 | 4.08 | 28.53* | 2.51 |
| AT 1000× 0.00075% vol. 1:150000 | 0.00 | 18.87 | 16.04 | 28.72 | 6.85 | 21.91 | 4.04 | 3.90 | 22.27* | 1.67 |

*In planta* evaluation supported the hypothesis that Atonik® treatment is efficient against *E. lata* pathogen. The inoculated *E. lata* treatment had significantly longer lesions than *Eutypa* + Atonik®, Atonik® alone, and the non-inoculated control. *Eutypa* + Atonik®, Atonik® alone, and the control were not significantly different from one another at p < 0.05 (Table 2). Fig. 1 demonstrates the graphical results of phytopathogenicity test and Fig. 2 shows the mean lesion length. The longest lesions were observed in grapevines inoculated with *E. lata* isolate E2 alone, with a mean lesion length of 48.38 mm. This treatment was assigned to LSD group “a” and differed significantly from all other treatments. In contrast, grapevines inoculated with *E. lata* and treated with Atonik® showed a markedly lower mean lesion length of 12.29 mm and were assigned to LSD group “b”. The non-inoculated control showed only limited necrosis, with a mean lesion length of 5.43 mm, while no lesions were observed in grapevines treated with Atonik® alone.

**Figure 1.**
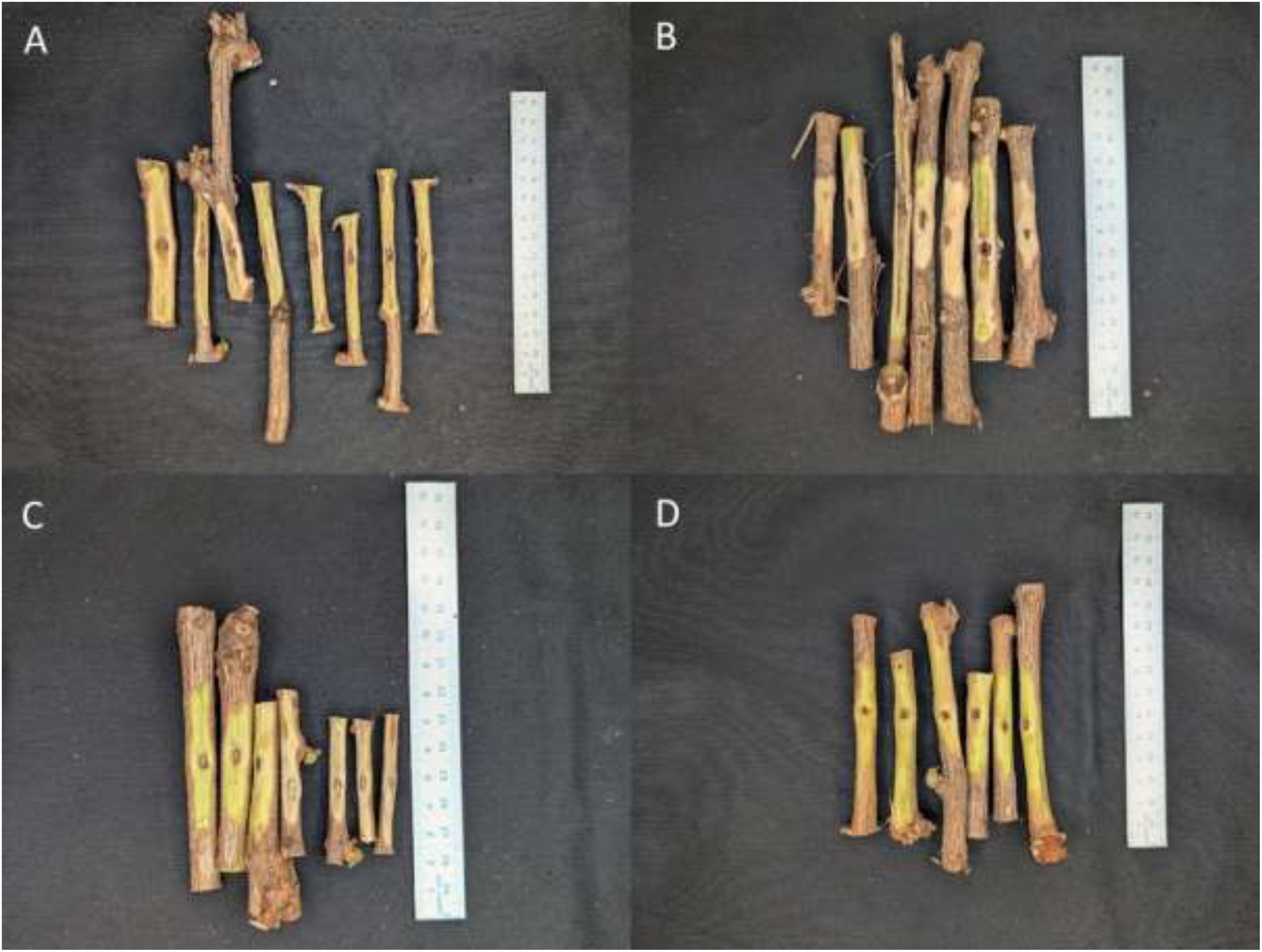
Pathogenicity test. (A) Lesions caused by *E. lata* strain E2; (B) lesions caused by *E. lata* strain E2 in grapevines treated with Atonik®; (C) non-inoculated control; (D) non-inoculated control treated with Atonik®.

**Figure 2.**
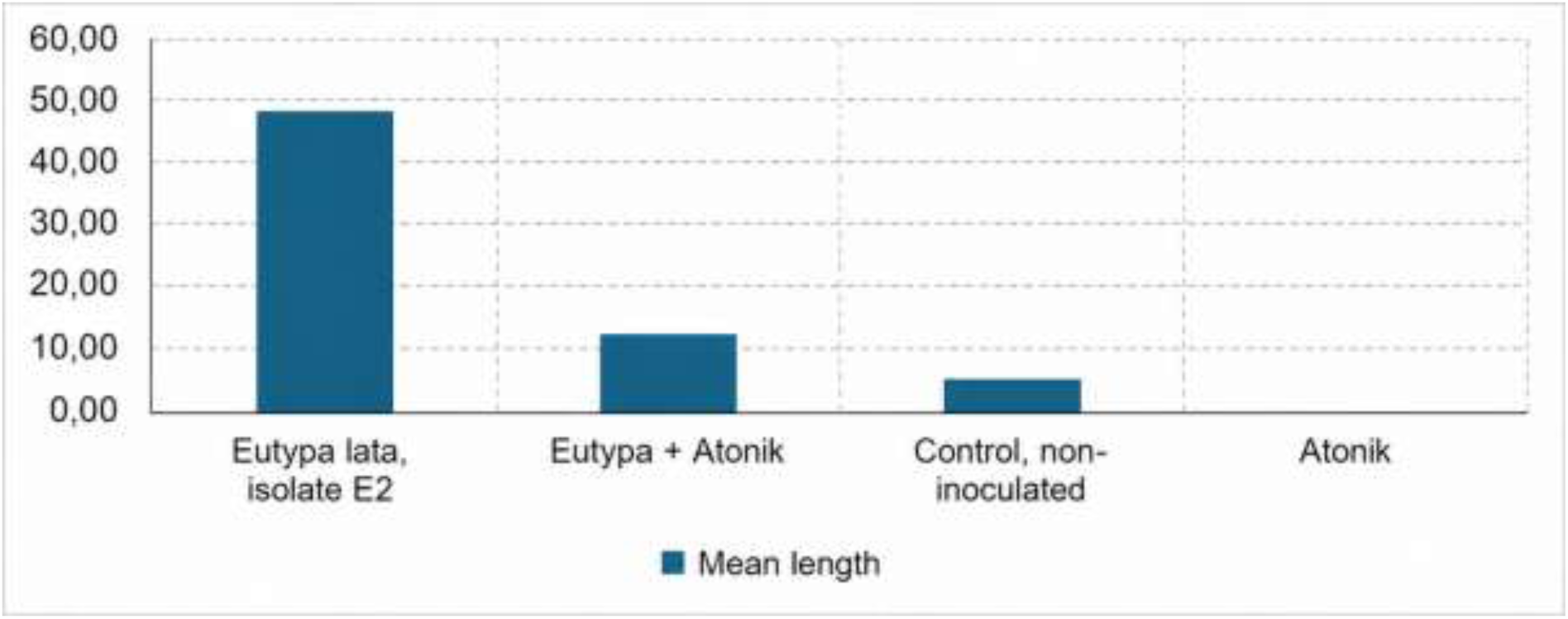
The highest mean lesion length was observed in plants inoculated with *E. lata* isolate E2 alone, whereas treatment with Atonik® markedly reduced lesion development in inoculated grapevines. No necrosis was observed in grapevines treated with Atonik® alone, while only limited lesion development was recorded in the non-inoculated control. Mean lengths are depicted in mm on axis y.

**Table 2.** Measured values of necrosis in grapevines inoculated with *Eutypa lata* isolate E2 and treated with Atonik®. One-way ANOVA was calculated across treatments. Fisher’s LSD pairwise comparisons were computed using the pooled error mean square from the ANOVA: LSD = t(0.975, df_error) × sqrt(MSE × (1/n1 + 1/n2)).

| Treatment | n | Mean (mm) | SD | SE | LSD group |
| --- | --- | --- | --- | --- | --- |
| *Eutypa lata*, isolate E2 (E8) | 8 | 48.38 | 11.94 | 4.22 | **a** |
| *Eutypa* + Atonik® (EA7) | 7 | 12.29 | 24.83 | 9.39 | **b** |
| Control, non-inoculated (K7) | 7 | 5.43 | 9.55 | 3.61 | **b** |
| Atonik® (A6) | 6 | 0.00 | 0.00 | 0.00 | **b** |

### Gene expression quantification of VvSOD gene

The expression of the *VvSOD* gene was evaluated by absolute qPCR quantification across four treatments: grapevines inoculated with *E. lata* isolate E2, grapevines inoculated with *E. lata* and treated with Atonik®, non-inoculated control grapevines, and grapevines treated with Atonik® alone. The standard curve generated from the synthesized *VvSOD* DNA standard was described by the equation Ct = −3.333 × log₁₀(concentration) + 13.852, with R² = 0.9625 and a PCR efficiency of 99.5% (Fig. 3).

**Figure 3.**
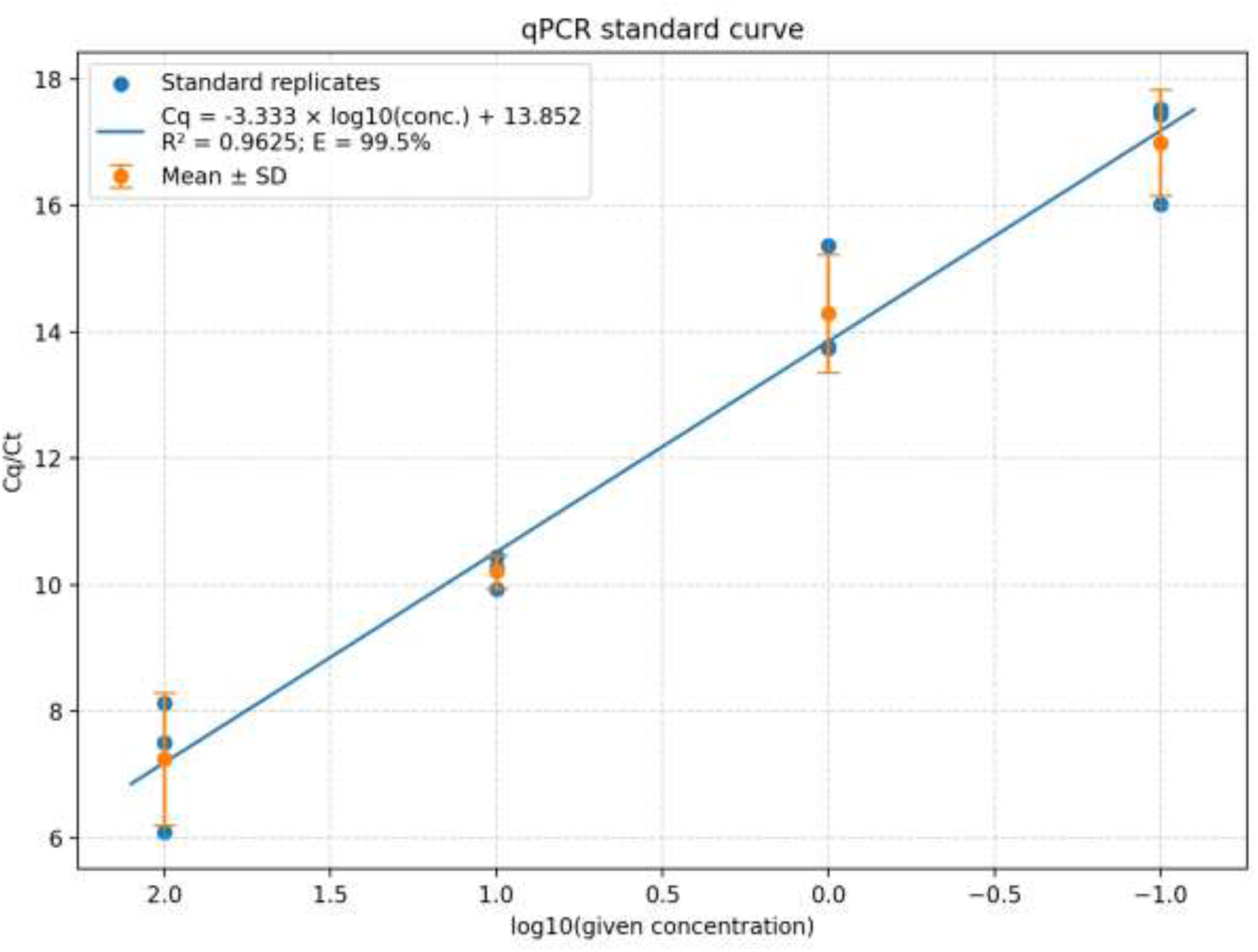
Standard curve for absolute qPCR quantification of the target *VvSOD* sequence. The curve was generated from a 10-fold serial dilution series of the synthesized DNA standard, corresponding to concentrations of 6 × 10^10^, 6 × 10^9^, 6 × 10^8^, and 6 × 10^7^. Ct values were estimated from raw amplification curves using a fluorescence threshold of 1.0. The resulting calibration curve was described by the equation Ct = −3.333 × log₁₀(concentration) + 13.852, with R² = 0.9625 and a PCR efficiency of 99.5%.

The highest detection rate was observed in grapevines inoculated with *E. lata* isolate E2, where 21 out of 24 technical replicates were positive, corresponding to a detection rate of 87.5%. Atonik® treated grapevines showed 13 positive reactions out of 18 technical replicates, corresponding to a detection rate of 72.2%. In the non-inoculated control, 14 out of 21 technical replicates were positive, giving a detection rate of 66.6%, whereas the *E. lata* + Atonik® treatment showed 13 positive reactions out of 21 technical replicates, corresponding to a detection rate of 61.9%.

Mean Ct values differed among treatments (Table 3). The lowest mean Ct value was recorded in the *E. lata* + Atonik® treatment (33.11; 998 copies.ng^-1^), followed by Atonik® alone (33.81; 616 copies.ng^-1^). In contrast, higher mean Ct values were observed in grapevines inoculated with *E. lata* isolate E2 alone (35.62; 177 copies.ng^-1^) and in the non-inoculated control (35.67; 171 copies.ng^-1^). Based on the standard curve, the estimated absolute quantities were 1.66 × 10⁻⁶ for the *E. lata* + Atonik® treatment, 1.03 × 10⁻⁶ for Atonik® alone, 2.94 × 10⁻⁷ for *E. lata* isolate E2, and 2.85 × 10⁻⁷ for the non-inoculated control.

**Table 3.** qPCR detection summary for *VvSOD* quantification across treatments. Mean Ct values were calculated only from positive technical replicates. Detection rate represents the proportion of positive technical replicates relative to the total number of reactions.

| Treatment | Bio samples | Positive tech reps | Total tech reps | Detection rate | Mean Ct | Abs. Quantification (copies.ng<sup>-1</sup>) |
| --- | --- | --- | --- | --- | --- | --- |
| *Eutypa lata*, isolate E2 (E8) | 8 | 21 | 24 | 0.875 | 35.62 | 177 |
| *Eutypa* + Atonik® (EA7) | 7 | 13 | 21 | 0.619 | 33.11 | 998 |
| Control, non-inoculated (K7) | 7 | 14 | 21 | 0.666 | 35.67 | 171 |
| Atonik® (A6) | 6 | 13 | 18 | 0.722 | 33.81 | 616 |

Overall, the lowest Ct values and the highest estimated absolute quantities of the *VvSOD* target were observed in treatments involving Atonik®, particularly in grapevines inoculated with *E. lata* and treated with Atonik®. This indicates an increased abundance of the *VvSOD* transcript in Atonik® treated grapevines compared with untreated inoculated and non-inoculated controls.

### Transcriptomic response of grapevine to Eutypa lata infection and Atonik® treatment

168 genes respond to *E. lata* isolate E2 infection in the wood padj < 0.05 and |log2FC| ≥1 and 43 genes respond to the Atonik® treatment using the same criteria. The original summary(res_infection) indicated 199 significant genes based solely on an adjusted p-value (padj) < 0.05. After applying the additional criterion of |log2FoldChange| ≥ 1, 168 genes remained.

Principal component analysis (PCA) revealed marked differences in the global transcriptomic profiles of the four samples (Fig. 4). PC1 explained 63% of the total variance and separated the control samples (A6 and K7) from the infected samples (EA7 and E8), whereas PC2 explained 32% of the variance and appeared to distinguish treated from untreated samples. Together, the first two principal components accounted for 95% of the total variance. PCA indicated that infection status was the dominant source of transcriptomic variation, accounting for much of the separation along PC1, whereas treatment primarily contributed to separation along PC2. This pattern suggests distinct and approximately additive effects of infection and treatment on the overall gene expression.

**Figure 4.**
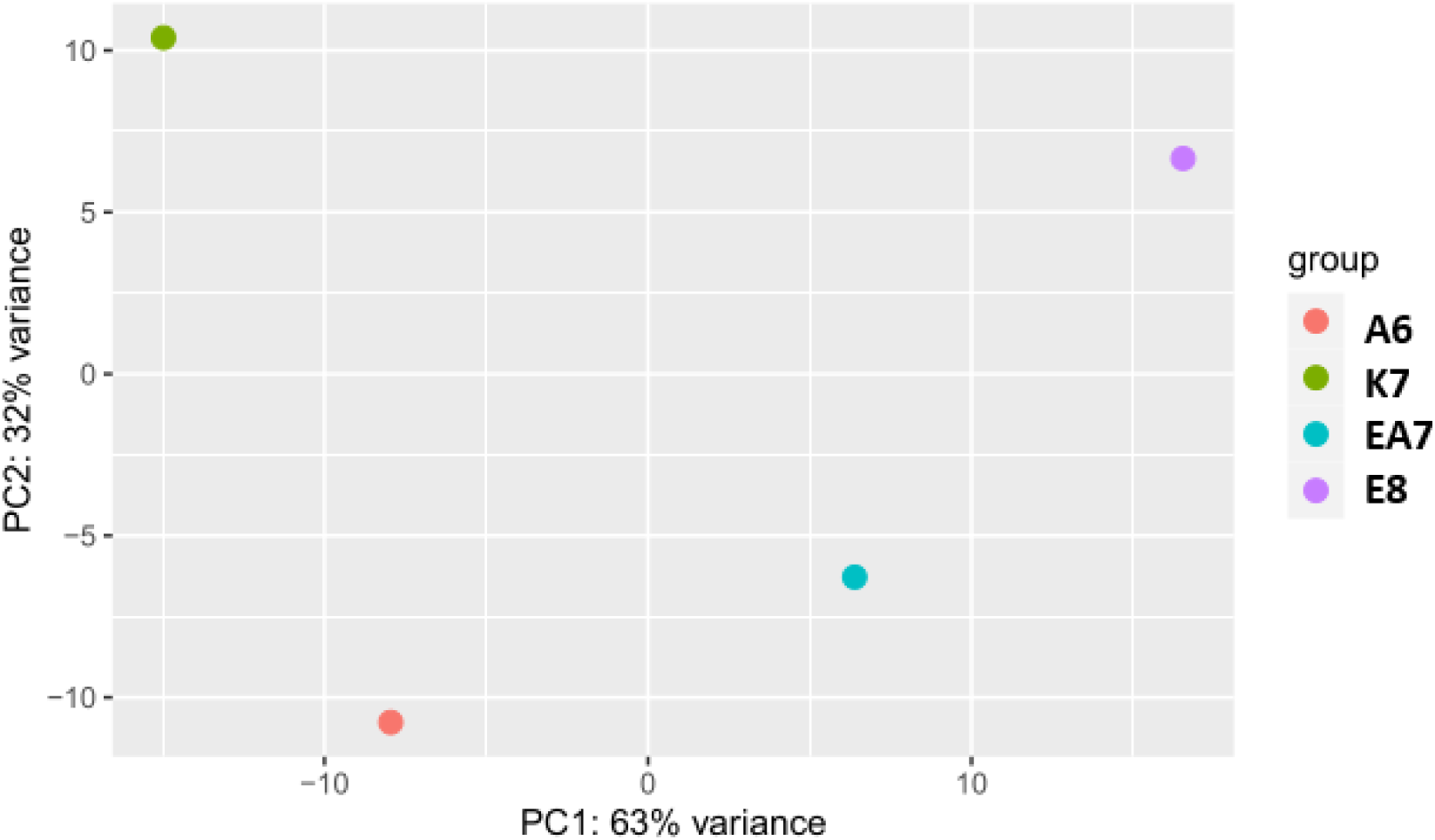
Principal component analysis of RNA-seq samples based on variance stabilized gene expression counts. PC1 and PC2 explained 63% and 32% of the total variance, respectively. Colours indicate the treatments.

The overlap of expressed genes among the RNA-seq libraries is shown in Figure 5. Using a threshold of DESeq2 normalized counts ≥ 10, a total of 12,441 genes were detected in at least one of the four RNA-seq libraries. The highest number of expressed genes was detected in A6 (11,548 genes), followed by EA7 (11,458 genes), E8 (11,193 genes), and K7 (10,621 genes). A large core of 9,156 genes, corresponding to 73.6% of all detected genes, was shared among all four libraries. Only small numbers of genes were detected exclusively in individual libraries: 13 in A6, 59 in EA7, 54 in E8, and 70 in K7. Among the pairwise overlaps, 305 genes were shared exclusively by the two Atonik® treated libraries A6 and EA7, whereas 166 genes were shared exclusively by the untreated libraries K7 and E8. Similarly, 323 genes were shared exclusively by the non-infected libraries A6 and K7, while 246 genes were shared exclusively by the infected libraries EA7 and E8. The largest three-library-specific overlap comprised 995 genes detected in A6, EA7, and E8 but not in K7. Overall, the analysis revealed a highly conserved set of expressed genes across the four libraries, accompanied by comparatively small library and condition associated subsets.

**Figure 5.**
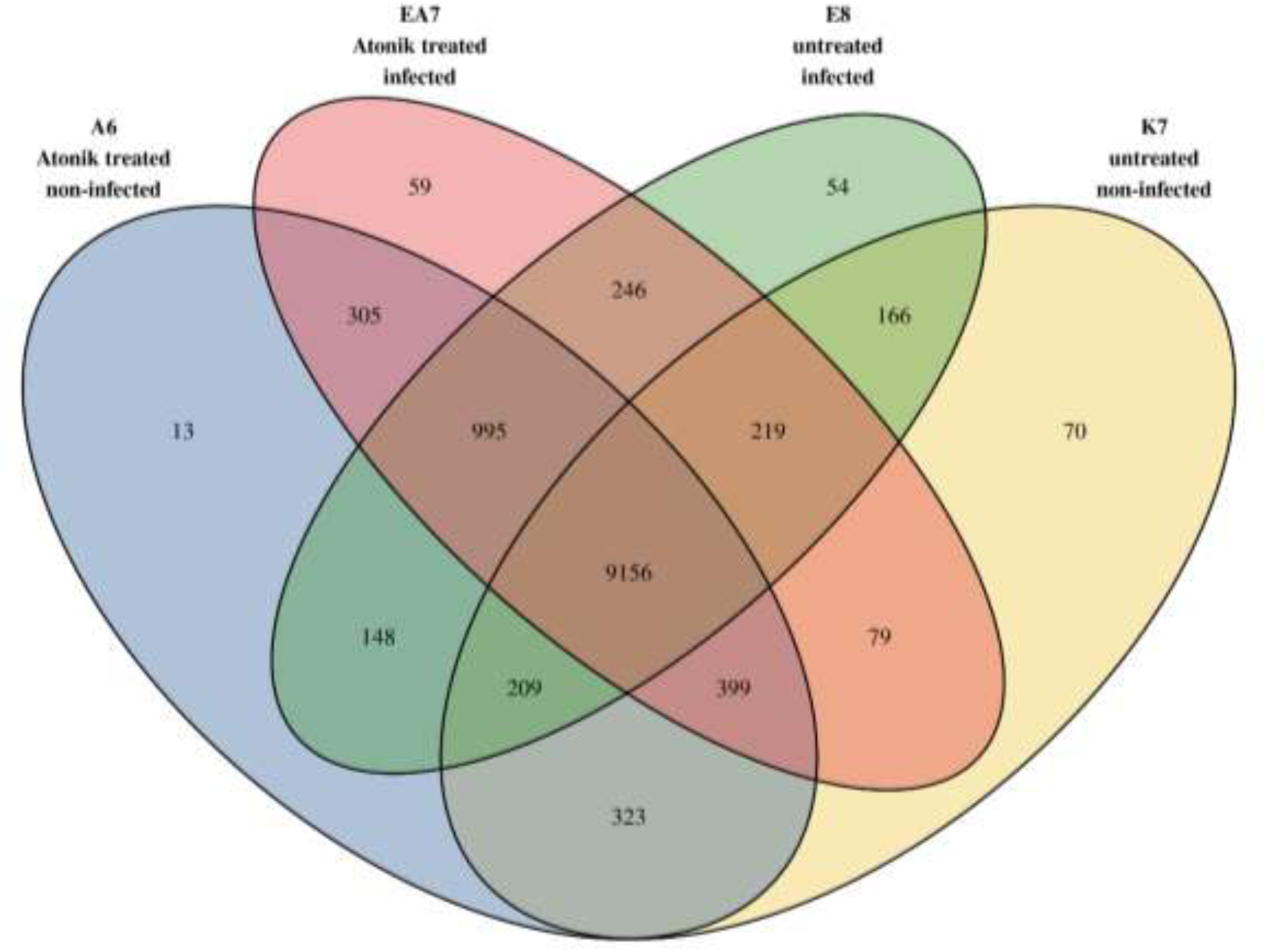
Overlap of expressed genes among the four RNA-seq libraries A6, K7, EA7 and E8. Genes with DESeq2 normalized counts ≥ 10 in a given library were considered expressed. The analysis is descriptive and does not represent differential expression testing among the individual libraries.

### Differential gene expression associated with E. lata isolate E2 infection and Atonik® treatment

The four RNA-seq libraries represented a 2 × 2 experimental arrangement: A6 corresponded to the Atonik® treated non-infected control, K7 to the untreated non-infected control, EA7 to the Atonik® treated infected sample, and E8 to the untreated infected sample. Differential expression was evaluated using an additive model including infection and treatment as independent factors. Thus, the estimated *E. lata* isolate E2 infection effect represented the common transcriptional difference between EA7 and A6 under treated conditions and between E8 and K7 under untreated conditions.

Using an adjusted p-value threshold of 0.05 and an absolute shrunken log2 fold-change of at least 1, 168 genes were associated with infection status. Of these, 116 genes showed higher expression in the infected samples EA7 and E8, whereas 52 genes showed higher expression in the non-infected controls A6 and K7 (Supplementary Table S1). The infection associated transcriptional response included genes related to pathogen defence and stress signalling, such as pathogenesis-related protein PR-4, acidic chitinase, chitinase 10, WRKY transcription factors, ethylene-responsive transcription factors and disease-resistance-associated receptor-like kinases.

Several genes associated with phenylpropanoid and specialized metabolism showed higher expression in the infected samples EA7 and E8, including multiple stilbene synthase 4-like genes, dirigent proteins, laccase 14, caffeic acid 3-O-methyltransferase, and cytochrome P450 71D9. In contrast, several genes encoding components of photosystems I and II, a ribulose-1,5-bisphosphate carboxylase/oxygenase small-subunit protein, and other chloroplast-associated proteins showed higher expression in the non-infected controls A6 and K7. This expression pattern suggests an infection associated shift towards defence related and specialized metabolic processes, accompanied by lower expression of selected photosynthesis related genes in the infected samples.

The most significant infection associated gene was LOC100253541, annotated as dirigent protein 22 (Supplementary material S3), which showed higher expression in EA7 and E8 than in A6 and K7 (log2FC = 3.29, adjusted p = 2.11 × 10⁻¹⁵). Other strongly induced genes included MLP-like protein 34 (LOC100267971; log2FC = 4.30), pathogenesis-related protein PR-4 (LOC100255405; log2FC = 3.17), WRKY transcription factor 55 (LOC100263918; log2FC = 4.17), and stilbene synthase 4 (LOC132252668; log2FC = 3.33) (Table 4).

**Table 4.** Most significant genes associated with *E. lata* isolate E2 infection in grapevine. The table presents selected genes with the strongest infection-associated transcriptional changes identified using the additive DESeq2 model ∼ infection + treatment. Positive log₂ fold-change values indicate higher expression in the infected samples EA7 and E8 relative to the non-infected controls A6 and K7. Genes were selected from the set meeting the criteria of adjusted *p* < 0.05 and |shrunken log₂FC| ≥ 1. The adjusted *p*-values were corrected for multiple testing using the Benjamini -Hochberg procedure.

| Gene ID | Annotation | log<sub>2</sub>FC<br>(Infected vs non-infected) | Adjusted p-value |
| --- | --- | --- | --- |
| LOC100253541 | Dirigent protein 22 | 3.29 | 2.11E-15 |
| LOC100267971 | MLP-like protein 34 | 4.30 | 6.35E-15 |
| LOC100255405 | Pathogenesis-related protein PR-4 | 3.17 | 1.22E-13 |
| LOC100263918 | WRKY transcription factor 55 | 4.17 | 3.05E-12 |
| LOC132252668 | Stilbene synthase 4 | 3.33 | 4.90E-12 |

### Treatment associated transcriptional responses in A6, EA7, K7 and E8

Differential expression was evaluated using an additive model containing infection status and treatment as independent factors. Thus, the estimated treatment effect represented the common transcriptional difference between A6 and K7 in the non-infected condition and between EA7 and E8 in the infected condition, under the assumption that the Atonik® treatment effect was comparable across infection states. Using an adjusted p-value threshold of 0.05 and an absolute shrunken log2 fold change of at least 1, 43 genes were associated with Atonik® treatment status. Of these, 33 genes showed higher expression in the A6 and EA7, whereas 10 genes showed higher expression in the untreated samples K7 and E8. The identified set comprised 41 protein-coding genes and two long non-coding RNAs; nine genes were annotated as uncharacterized (Supplementary Table S2).

Genes showing higher expression in A6 and EA7 included several genes potentially related to plant defence and stress responses, including an osmotin-like protein, defensin Ec-AMP-D2, snakin-1, snakin-2, putative lipid-transfer protein DIR1, NDR1/HIN1-like protein 1, peroxidase 4, an ethylene-responsive transcription factor TINY, and two small heat-shock proteins. Increased expression was also observed for genes related to cell-wall organisation and modification, including proline-rich cell-wall proteins, non-classical arabinogalactan protein 31, pectin acetylesterase 8, pectinesterase, and IRX15-LIKE (Supplementary Table S2).

In contrast, genes showing higher expression in the untreated samples K7 and E8 included pathogenesis-related protein PR-4, dirigent protein 22, caffeic acid 3-O-methyltransferase, loganic acid O-methyltransferase, and 1-aminocyclopropane-1-carboxylate oxidase homolog 1. The occurrence of defence associated genes in both expression directions indicates that the treatment was associated with transcriptional reprogramming of multiple defence and stress related pathways rather than with a uniform induction of the entire defence response.

The most significant treatment associated transcript was the uncharacterized long non-coding RNA LOC100257577, which showed higher expression in A6 and EA7 than in K7 and E8 (log2FC = 2.82, adjusted p = 3.14 × 10⁻¹¹). Among the annotated genes, a 14 kDa proline-rich protein DC2.15 (LOC100267804; log2FC = 3.51), cysteine proteinase inhibitor 8-like (LOC132253257; log2FC = 2.82), osmotin-like protein (LOC100253492; log2FC = 2.28), and defensin Ec-AMP-D2 (LOC100242603; log2FC = 1.93) showed higher expression in the treated samples. Cytochrome P450 78A3 (LOC100258159; log2FC = 4.69) and galactinol-sucrose galactosyltransferase (LOC100251818; log2FC = 4.53) displayed the largest positive expression differences. Conversely, pathogenesis-related protein PR-4 (LOC100255405) showed markedly higher expression in the untreated samples K7 and E8 (log2FC = −3.51, adjusted p = 1.01 × 10⁻⁹). Dirigent protein 22 (LOC100248431; log2FC = −2.48), caffeic acid 3-O-methyltransferase (LOC100232921; log2FC = −1.40), and 1-aminocyclopropane-1-carboxylate oxidase homolog 1 (LOC104879286; log2FC = −1.23) were also more highly expressed in the untreated samples (Table 5, Supplementary Table S2).

**Table 5.** Most significant transcripts associated with Atonik® treatment in grapevine. The table lists selected genes showing higher expression in the Atonik® treated samples A6 and EA7 than in the untreated samples K7 and E8. Differential expression was evaluated using an additive DESeq2 model including infection status and treatment as independent factors. Positive shrunken log₂ fold-change values indicate higher expression in the treated samples. Genes were selected using the thresholds adjusted p < 0.05 and |shrunken log₂FC| ≥ 1. Adjusted p-values were corrected for multiple testing using the Benjamini-Hochberg procedure.

| Gene ID | Annotation | log<sub>2</sub>FC<br>(Treated vs<br>untreated) | Adjusted p-value |
| --- | --- | --- | --- |
| LOC100257577 | uncharacterized LOC100257577 | 2.82 | 3.14E-11 |
| LOC100267804 | 14 kDa proline-rich protein DC2.15 | 3.51 | 6.13E-08 |
| LOC132253257 | cysteine proteinase inhibitor 8-like | 2.82 | 2.73E-07 |
| LOC100253492 | osmotin-like protein | 2.28 | 4.81E-06 |
| LOC100242603 | defensin Ec-AMP-D2 | 1.93 | 6.76E-06 |
| LOC100258159 | cytochrome P450 78A3 | 4.69 | 7.38E-06 |
| LOC100251818 | galactinol--sucrose galactosyltransferase | 4.53 | 1.06E-05 |

No transcript annotated as superoxide dismutase met the applied significance criteria for the treatment associated contrast (Supplementary table S4). However, Peroxidase 4 (LOC100257005) showed higher expression in the treated group represented by A6 and EA7 than in the untreated group represented by K7 and E8 (log2FC = 1.91, adjusted p = 0.0197). This result suggests Atonik® treatment associated modulation of reactive oxygen species metabolism, although specific induction of SOD was not demonstrated by RNA-seq (Supplementary table S2).

## Discussion

Fungal trunk diseases represent one of the most challenging disease complexes affecting grapevine and other perennial woody crops ^1^. Their management is complicated by the diversity of associated pathogens, the long latent phase of infection, the progressive colonization of woody tissues, and the limited availability of curative treatments ^2,20^. For this reason, current control strategies rely mainly on preventive approaches, including the use of healthy planting material, pruning wound protection, sanitation practices, biological control agents, and compounds capable of reducing pathogen development or stimulating host defence responses ^10^. In this context, sodium nitrophenolates represent an interesting group of plant growth regulator and stimulant like compounds that may affect plant physiology, stress responses, and defence related metabolism.

### In vitro antifungal activity of sodium nitrophenolates

The *in vitro* assays showed that sodium nitrophenolate based treatments affected the mycelial growth of selected fungal trunk disease associated pathogens, although the response differed among fungal species and treatment concentrations. Regarding the used spectra of the used dilutions Atonik® showed the most persistent activity across dilutions against *B. dothidea* BD (2-1) even at concentration 0.00075% and the highest percentage levels reached the efficiency against *C. luteo-olivacea* CLO (H20), 74.62 % of inhibition. The lowest efficiency was observed against *F. mediterranea* FM (FOP1), 25.62 % of inhibition and *D. torresensis* DT (CZ2), 33.13 % of inhibition by 0.75% solution of Atonik®. Although Atonik® and sodium nitrophenolate based formulations have rarely been evaluated as direct antifungal agents, previous *in vitro* studies have tested Atonik® on agar media against selected phytopathogenic fungi using Petri dish based assays. Djanaguiraman et al. ^21^ evaluated the compatibility of Atonik® with fungicides using a poisoned food technique and measured radial mycelial growth of *Fusarium oxysporum* f. sp. *vasinfectum*, *Colletotrichum capsici*, *Alternaria alternata*, and *Botrytis cinerea*, conclusion of this study was that Atonik® was compatible with carbendazim a fytolon. Similarly,

Manole and Iacomi ^22^ tested Atonik® on PDA medium against *Alternaria* spp. pathogenic to tomato. Manole and Iacomi ^22^ reported that Atonik® at 0.05% inhibited the *in vitro* mycelial growth of selected *Alternaria* species pathogenic to tomato, including *A. solani*, *A. lycopersici*, and *A. tenuissima*. However, Atonik® did not inhibit spore germination of the tested *Alternaria* species, indicating that its antifungal effect was limited to mycelial growth and was dependent on the developmental stage of the pathogen. However, to our knowledge, comparable *in vitro* assays against grapevine trunk disease associated fungi have not been reported.

Similar concentration and compound dependent effects have been reported for natural phenolic compounds tested against GTD associated fungi, where eugenol, thymol and EGCG differed markedly in their ability to inhibit mycelial growth, spore germination and infection capacity ^23,24^. Recent evidence also indicates that compound sodium nitrophenolate can directly inhibit mycelial growth of some phytopathogenic fungi, although this has so far been shown outside the GTD pathosystem ^25^. This indicates that the tested compounds do not act as a broad spectrum fungicide with uniform activity against all pathogens, but rather exert species specific effects that may depend on fungal physiology, growth rate, and sensitivity to phenolic or nitrophenolic compounds. Such variability is expected within the GTD pathosystem, which comprises taxonomically and ecologically diverse fungi with different colonization strategies and stress tolerance mechanisms ^8,12,20^. The observed inhibition of selected pathogens suggests that sodium nitrophenolates may contribute to the suppression of fungal growth under controlled conditions. However, the *in vitro* effect should be interpreted with caution, as direct inhibition on artificial media does not necessarily reflect efficacy *in planta*, where compound uptake, tissue distribution, host metabolism, and plant pathogen interactions can strongly influence the final outcome ^12,23,26^. Importantly, sodium nitrophenolates are primarily described as plant growth regulator or stimulant like compounds rather than conventional fungicides, which is consistent with their regulatory classification as nitrophenolate plant growth regulators ^27,28^. Therefore, their potential role in trunk disease management may not be limited to direct inhibition of pathogen growth. Their effect may also involve modulation of host physiology and stress related defence responses, including antioxidant metabolism, as previously reported for nitrophenolate based stimulants and grapevine defence stimulating compounds ^9,14^. This dual perspective is particularly relevant for grapevine trunk diseases, for which effective management remains difficult and relies mainly on integrated preventive strategies and treatments capable of reducing pathogen development or enhancing host defence capacity ^2,12^.

Overall, the *in vitro* results support the hypothesis that sodium nitrophenolates may influence the development of fungal trunk disease associated pathogens, but their efficacy is likely pathogen dependent and concentration dependent. The *in vitro* assays provided preliminary evidence of antifungal activity, but *in planta* validation was required to confirm their biological relevance.

### In planta effect of Atonik® on Eutypa lata induced lesion development

In the present study, the *in planta* effect of sodium nitrophenolates was evaluated against *E. lata*, one of the most important causal agents of grapevine trunk diseases ^2,12,29^. The *in planta* experiment demonstrated that inoculation with *E. lata* isolate E2 caused extensive lesion development in grapevine wood, confirming the pathogenic potential of this isolate under experimental conditions. Grapevines inoculated with *E. lata* alone developed the longest lesions, with a mean lesion length of 48.38 mm, and this treatment differed significantly from all other treatments. In contrast, grapevines inoculated with *E. lata* and treated with Atonik® showed a markedly lower mean lesion length of 12.29 mm. This reduction suggests that Atonik® treatment may limit lesion development caused by *E. lata* or may enhance the ability of grapevine tissues to restrict pathogen colonization.

Interestingly, the lesion length observed in the *E. lata* + Atonik® treatment did not differ significantly from the non-inoculated control or from gprapevines treated with Atonik® alone. This result indicates that the application of Atonik® substantially reduced disease expression under the conditions of the experiment. However, the high variability observed in the *E. lata* + Atonik® treatment, as reflected by the relatively large standard deviation, suggests that the response to treatment was not uniform among individual grapevines. Such variability is common in woody plant pathosystems, where lesion development and symptom expression can be influenced by host genotype, plant physiological status, wound response, local tissue conditions, environmental stress, and the establishment success of the pathogen ^20,29,30^.

The absence of lesions in grapevines treated with Atonik® alone indicates that the tested formulation did not induce visible phytotoxic necrosis under the experimental conditions. Compounds considered for use in trunk disease management should not negatively affect host tissues, particularly when applied to stressed or wounded grapevines. The limited necrosis observed in the non-inoculated control most likely reflects a wound response associated with the inoculation procedure rather than pathogen driven lesion development. Together, these results support the hypothesis that sodium nitrophenolate based treatments may contribute to reducing wood necrosis caused by GTD associated pathogens.

### Effect of Atonik® on VvSOD expression during Eutypa lata infection

The analysis of *VvSOD* expression suggests that Atonik® treatment may influence antioxidant related responses in grapevine. Superoxide dismutase is a key enzyme involved in the detoxification of reactive oxygen species, which play an important role in plant responses to biotic and abiotic stress ^31,32^. In the present study, the lowest mean Ct values and the highest estimated *VvSOD* abundance were observed in Atonik® treated grapevines, particularly in grapevines inoculated with *E. lata* and treated with Atonik® (mean Ct 33.1). This trend corresponded with the reduced lesion development observed in the *in planta* experiment, suggesting that sodium nitrophenolates may contribute to limiting disease expression through modulation of host stress responses. This interpretation is consistent with previous reports describing Atonik® and nitrophenolate based products as compounds capable of affecting plant physiology, metabolism, and stress related responses ^14^.

In contrast, grapevines inoculated with *E. lata* alone and non-inoculated control grapevines showed higher Ct values and lower estimated *VvSOD* abundance (mean Ct 35.6 and 35.7).

### Transcriptomic response of grapevine to Eutypa lata infection and Atonik® treatment

RNA-seq analysis indicated that *E. lata* infection was the main driver of transcriptomic variation in grapevine wood. This was supported by PCA, where infected samples separated from non-infected controls mainly along PC1, and by differential expression analysis, which identified 168 infection associated genes. The infection response was characterized by higher expression of genes related to defence signalling and pathogen response, including PR-4, chitinases, WRKY transcription factors, ethylene-responsive transcription factors and receptor-like kinases. Similar activation of PR proteins, chitinases, WRKY factors and other defence related genes has previously been reported in grapevine tissues challenged by *E. lata* and other GTD-associated fungi ^33,34^.

Infected grapevines also showed increased expression of genes involved in phenylpropanoid and specialized metabolism, including stilbene synthase genes, dirigent proteins, laccase 14 and caffeic acid 3-O-methyltransferase. This is consistent with previous reports showing that grapevine responses *to E. lata* involve activation of phenylpropanoid, flavonoid and stilbenoid pathways, with stilbenes representing important grapevine phytoalexins ^34,35^. In contrast, several photosynthesis and chloroplast associated genes showed higher expression in non-infected controls, suggesting a shift from primary metabolism towards defence related processes during infection.

Atonik® treatment was associated with a smaller but distinct transcriptional response, represented by 43 differentially expressed genes. Genes showing higher expression in Atonik® treated samples included osmotin-like protein, defensin Ec-AMP-D2, snakin-1, snakin-2, DIR1-like lipid-transfer protein, NDR1/HIN1-like protein 1, peroxidase 4, ethylene-responsive transcription factor TINY and small heat-shock proteins. This pattern suggests modulation of stress related, antimicrobial, ROS associated and cell wall related pathways. Such an interpretation is consistent with the reported mode of action of nitrophenolate based products, including Atonik®, which have been described as compounds affecting plant physiology, metabolism and stress responses rather than acting as conventional fungicides ^14^.

No transcript annotated as superoxide dismutase met the applied significance criteria for the treatment associated contrast. However, peroxidase 4 was more highly expressed in Atonik® treated samples, suggesting that Atonik® may influence reactive oxygen species metabolism through antioxidant pathways other than SOD. Plant peroxidases are directly involved in ROS metabolism, lignification, cell-wall reinforcement, and defence responses during plant pathogen interactions ^36,37^. Overall, the RNA-seq data support the hypothesis that Atonik® treatment may contribute to reduced disease expression by modulating host defence and stress related pathways, which is consistent with the reported effects of nitrophenolate based products on plant physiology and stress responses ^14^.

## Conclusions

This study showed that sodium nitrophenolate based treatment with Atonik® can influence both fungal growth and grapevine responses to trunk disease associated pathogen like *E. lata*. *In vitro* assays demonstrated pathogen and concentration dependent inhibition of mycelial growth, indicating that the antifungal activity of Atonik® is not uniform across fungal species. *In planta*, Atonik® markedly reduced lesion development caused by *E. lata* isolate E2 and did not induce visible phytotoxic symptoms when applied alone. Gene expression and RNA-seq analyses further suggested that Atonik® may contribute to disease reduction through modulation of host stress and defence related pathways, including antioxidant metabolism, antimicrobial proteins, cell wall associated responses, and phenylpropanoid related processes. Further studies with additional biological replicates and targeted validation of candidate defence genes are needed to clarify the underlying mode of action.

## Materials and Methods

### Chemical substances

The tested product was Atonik® (Asahi Chemical Europe s.r.o., Prague, Czech Republic). Atonik® is officially described as a plant growth regulator ^27,28^ and stimulant like preparation based on a mixture of three sodium nitrophenolates. According to the official product information, Atonik® contains sodium p-nitrophenolate, sodium o-nitrophenolate, and sodium 5-nitroguaiacolate (Fig. 6) at concentrations of 3 g.L^-1^, 2 g.L^-1^, and 1 g.L^-1^, respectively (Asahi Chemical Europe s.r.o.). This composition is also consistent with the regulatory documentation for Atonik®, which lists the active substances as sodium para-nitrophenolate, sodium ortho-nitrophenolate, and sodium 5-nitroguaiacolate in the same concentration ratio. Together, these active components form a 6 g.L^-1^ mixture of sodium nitrophenolates that is reported to affect basic physiological and metabolic processes in plants ^14^.

**Figure 6.**
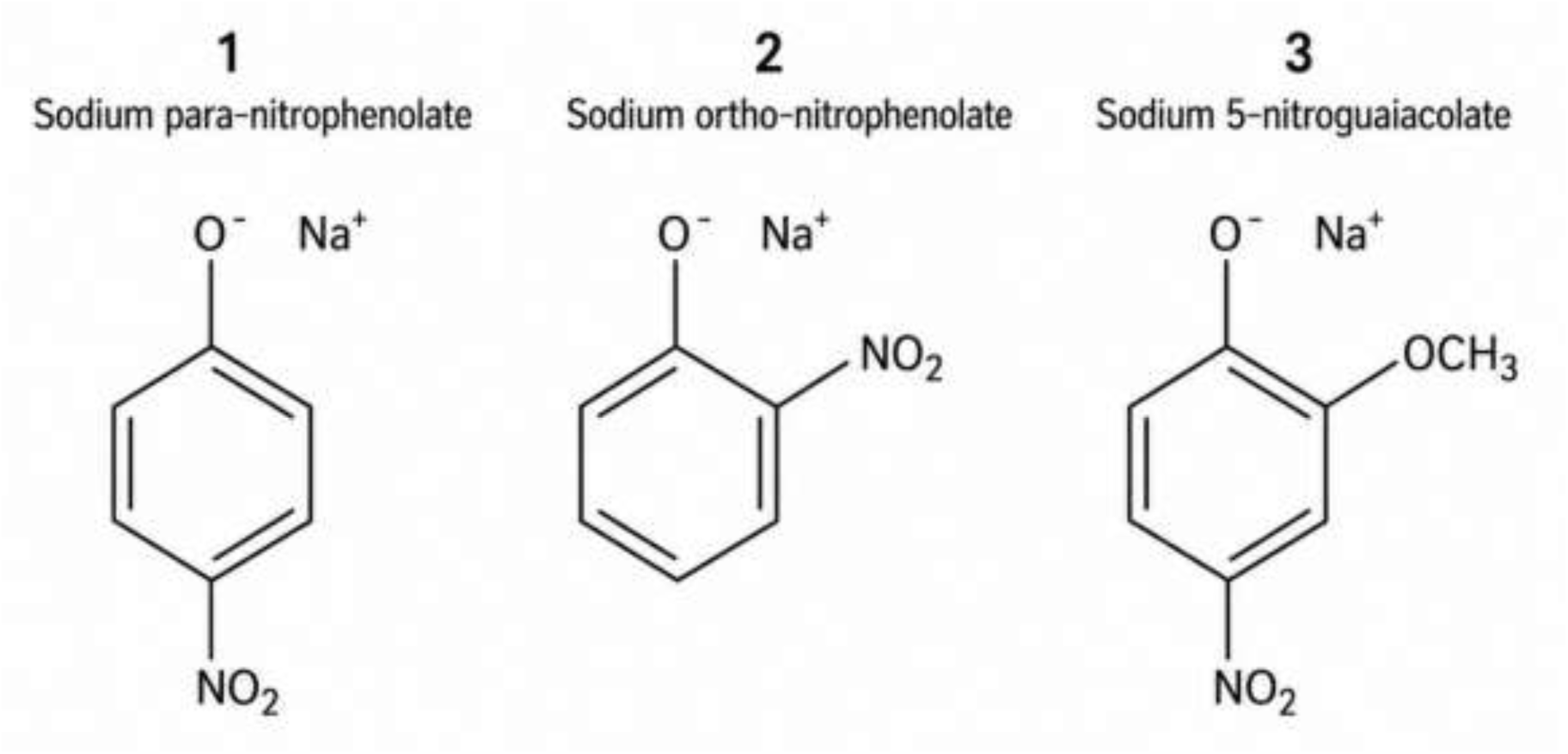
Chemical structures of the tested phenolates present in the Atonik® preparation.

### Fungal isolates

In vitro experiment used these isolates from Mendeleum collection: *Dactylonectria torresensis* (DT) MEND-F-0016, isolate CZ2 ^38^; *Cadophora luteo-olivacea* (CLO) MEND-F-0454, isolate H20 ^39^; *Eutypa lata* (EL) MEND-F-1156, isolate CS_12_27_4_KO ^40,41^; *Phaeoacremonium minimum* (PM) MEND-F-0013, isolate ESP1 ^42^; *Phaeomoniella chlamydospora* (PC) MEND-F-0229, isolate CS_61/14/2_ZN_H55 ^40^; *Diaporthe ampelina* (DA) MEND-F-157, isolate H23 ^42^; *Fomitiporia mediterranea* (FM) MEND-F-0471, isolate FOP1; *Diplodia seriata* (DS) MEND-F-0014 ^42^; *Botryosphaeria dothidea* (BD) MEND-F-0196, isolate 2-1; *Neofabraea kienholzii* (NK) MEND-F-1192 ^40^. *In planta* experiment used *Eutypa lata* isolate E2 (MEND-F-0403), described in this study, Acc. Nos. PZ564299 (beta-tubulin gene, *Tub2*) and PZ578101 (RNA polymerase II subunit 2 (RPB2) gene).

### Used concentrations of Atonik® for in vitro experiment

The tested substance of Atonik® was applied at four concentrations: 0.75% (v/v), corresponding to the stock working concentration, and three subsequent ten-fold dilutions, namely 10× at 0.075% (v/v), 100× at 0.0075% (v/v), and 1000× at 0.00075% (v/v). Substances were applied to PDA agar plates using a pipette (100 µl) and spread with an L-loop. A pathogen disk was transferred to the center of each dish. A positive control was also established. All treatments were established in triplicate. Mycelial growth was measured daily for 10 days at four positions per plate. The resulting data were processed using statistical software Statistica 14 CZ (StatSoft, Prague, Czech Republic).

### Atonik® efficacy in planta against Eutypa lata

Atonik® is registered in the Czech Republic for foliar application at a dose of 1 L ha⁻¹. The *in planta* experiment was carried out in 2025 in a net house at Mendeleum – Institute of Genetics, Mendel University in Brno, Lednice, Czech Republic. The experiment was performed using three year old cv.Sauvignon Blanc grapevines grafted onto SO4 rootstock. This cultivar was selected because of its high susceptibility to grapevine trunk disease associated pathogens ^29^. The grapevines had been grown for three years at the Mendeleum net house prior to inoculation. Inoculation with the E2 isolate was performed on June 6, 2025. Fifteen grapevines were inoculated with the selected highly aggressive isolate E2. For this study, the isolate was retrieved from storage and cultured on potato dextrose agar PDA (HiMedia, Mumbai, India) in Petri dishes incubated in the dark at 25 °C. After five days, mycelial plugs were taken from the growing margin of the colony and used to inoculate grapevine cuttings. Inoculations were performed according to the protocol described by Péros and Berger^43^. For the experiment, seven control grapevines were inoculated with PDA plugs without fungal mycelium (K7), and six additional control grapevines were treated in the same way but sprayed with Atonik® (A6). The first spray treatment was performed on August 4, 2025, using Atonik® at a concentration of 1.0%. The second treatment was applied on August 25, 2025, using Atonik® at 0.1%, and the third treatment was applied on September 15, 2025, also using Atonik® at 0.1%. Grapevines were subsequently harvested on October 7, 2025, for the measurement of lesion length and further analyses. Atonik® was diluted in tap water and applied to all green parts of the grapevines using a standard garden sprayer.

### Lesion measurement

After excavation, the roots were separated from the individual grapevines. Only the trunk sections containing either the inoculation site or the control wound were transported to the laboratory. In the laboratory, the inoculation site was carefully cleaned with a sterile scalpel, and lesions in the vascular tissues, particularly in the phloem, were measured. Detailed photographs were also taken for documentation (Fig. 1).

### Reisolation of Eutypa lata

For fungal re-isolation, necrotic wood tissue was first surface disinfected in 1% sodium hypochlorite for 3 min, followed by three rinses in sterile distilled water. The disinfected tissue was then cut into small fragments with a sterile scalpel. The wood pieces were aseptically placed on PDA amended with streptomycin (HiMedia, Mumbai, India) at 100 mg.L^-1^ to suppress bacterial contamination. Fungal colonies developing from the wood fragments were transferred to fresh PDA plates to obtain pure cultures. The obtained isolates were initially separated into a morphological groups corresponding to *Eutypa* like colonies. One representative isolate from each morphological group was selected for molecular confirmation. Mycelium from these pure cultures was harvested and used for genomic DNA extraction with the NucleoSpin Tissue Kit (Macherey-Nagel, Düren, Germany), according to the manufacturer’s instructions.

For molecular identification, *E. lata* was identified using *tub2* sequence data following Eichmeier et al. ^44^. PCR products were prepared for sequencing using the Mix2Seq Kit (Eurofins Genomics, Ebersberg, Germany), and Sanger sequencing was performed by Eurofins Genomics. The resulting sequences were compared with reference sequences deposited in GenBank using the BLAST algorithm to determine the identity of the re-isolated fungi ^45^.

### Extraction of RNA

RNA was extracted directly from 1 g of scraped phloem using the Spectrum Plant Total RNA Kit (Sigma-Aldrich, St. Louis, MO, USA). RNA extraction was done from all 26 grapevines, from the wood close to the inoculation point. Concentration was measured by SPECTROstar Nano. RNA was diluted to 5 ng.µl^-1^ using sterile nuclease-free water (Ambion, Pleasanton, California, USA).

### VvSOD gene expression analysis

Gene expression analysis was performed using the total RNA. Since *VvSOD* encodes superoxide dismutase, one of the key enzymes involved in antioxidant defence, the quantification was focused on the expression of this gene. This is a typical conserved motif of Fe/Mn-superoxide dismutases. This motif is described as a metal binding motif in Fe/Mn-SOD proteins ^46,47^. *VvSOD* expression was evaluated using real-time PCR. Primer sequences are VvFSD1 fwd (5′-GGACGACATTCCCATCATCA-3′) and rev (5′-GCAGCATTCCAAGACACAAG-3′) ^47^. Reverse transcription was performed following the protocol described by Eichmeier et al. ^48^, using the RevertAid RT Reverse Transcription Kit (Thermo Fisher Scientific, Waltham, Massachusetts, USA), resulting in complementary DNA (cDNA) samples. qPCR was carried out using Genaxxon mastermix (Genaxxon Bioscience GmbH, Ulm, Germany) on a RotorGene3000 (Corbett Research in Sydney, Australia). For absolute quantification of *VvSOD* expression, a synthesized DNA standard corresponding to the target amplicon was used. Absolute quantification was performed using a synthesized DNA standard corresponding to the target *VvSOD* amplicon (5′- ACATTCCCATCATCAGCTTGGACGTGTGGGAGCATGCTTATTATCTGGATTACAAGAACGACAAAGGGAAGTACGCGAATGTGTTCATGAACCACCTTGTGTCTTGGAAT-3′). Standard was delivered as 100 µM solution (according to formula 100 µM = 6.022 × 1013 copies/µL). A 10-fold serial dilution series of the standard was prepared and used to generate a calibration curve, in accordance with standard qPCR absolute quantification approaches and MIQE reporting recommendations ^49,50,51^.

### mRNA Library Preparation and Sequencing

For mRNA libraries was pooled four samples E8 (eight RNA samples: *Eutypa lata*, isolate E2), EA7 (seven RNA samples: *Eutypa* + Atonik®), K7 (seven RNA samples: control, non-inoculated) and A6 (six RNA samples: Atonik®). RNA from all samples used in the analysis was diluted to a concentration of 5 ng.µl^-1^ using sterile nuclease-free water (Ambion, Pleasanton, California, USA).

RNA sequencing libraries were prepared using the NEBNext Single Cell/Low Input RNA Library Prep Kit for Illumina (New England Biolabs, Ipswich, MA, USA) according to the manufacturer’s instructions. Individual libraries were indexed using NEBNext Multiplex Oligos for Illumina with Unique Dual Index Primer Pairs (New England Biolabs). The quality and fragment size distribution of the prepared libraries were assessed using an Agilent 2100 Bioanalyzer with the High Sensitivity DNA Kit (Agilent Technologies, Santa Clara, CA, USA). Library concentrations were determined using the Collibri Library Quantification Kit (Invitrogen, Thermo Fisher Scientific, Waltham, MA, USA) and the Qubit 1× dsDNA High-Sensitivity Assay Kit (Invitrogen, Thermo Fisher Scientific). Library integrity and size distribution were additionally verified by electrophoresis using 2% E-Gel EX agarose gels (Invitrogen, Thermo Fisher Scientific). The indexed libraries were sequenced on a NovaSeq X Plus platform (Illumina, San Diego, CA, USA), generating 150-bp paired-end reads (2 × 150 bp).

### Quality control and preprocessing of sequencing reads

The quality of the raw paired-end FASTQ files was assessed individually using FastQC v0.12.1 ^52^. The evaluated parameters included the total number of reads, per-base sequence quality, GC-content distribution, sequence duplication levels, adapter content, per-base sequence composition, and the presence of overrepresented sequences potentially corresponding to adapters, ribosomal RNA, or highly abundant transcripts. Adapter sequences and low-quality terminal bases were removed using fastp v0.23.4 ^53^, with automatic adapter detection for paired-end reads (--detect_adapter_for_pe) and quality-based trimming from the read tails (--cut_tail). After trimming, the processed FASTQ files were reassessed using FastQC to verify the quality of the retained reads and the effectiveness of preprocessing.

### Read alignment, gene-level quantification, differential expression analysis, and functional

Following quality control and preprocessing, the trimmed paired-end reads were aligned to the *Vitis vinifera* reference genome assembly GCF_030704535.1 (ASM3070453v1) using STAR v2.7.10b ^54^. The genome index was generated from the corresponding reference genome sequence and the NCBI RefSeq annotation release GCF_030704535.1-RS_2023_09. Paired-end reads were supplied to STAR as mate pairs. The resulting BAM files were coordinate sorted, indexed, and checked using SAMtools ^55^. Gene level read counts were generated from the coordinate sorted BAM files using featureCounts, implemented in the Subread package ^56,57^, together with the corresponding annotation file Vitis_vinifera_GCF_030704535.1.gtf. Paired-end fragments were assigned to annotated genomic features and summarized using the gene_id attribute, with library strandedness specified according to the library preparation protocol. Raw, unnormalized integer counts were combined into a count matrix containing 28,162 genes across four RNA-seq libraries. The four libraries represented a 2 × 2 experimental arrangement combining infection status and Atonik® treatment. A6 represented the Atonik® treated non-infected control, K7 the untreated non-infected control, EA7 the Atonik® treated sample infected with *E. lata* isolate E2, and E8 the untreated infected sample. Differential gene expression analysis was performed in R version 4.3.3^58^ using DESeq2 ^59^. Raw gene level counts were supplied as integers to DESeqDataSetFromMatrix(). Genes were retained when they had at least 10 reads in at least two libraries ^59^. After this filtering step, 12,449 genes remained for downstream analysis. An additive model containing infection status and treatment was fitted using the design formula: ∼ infection + treatment. The model estimated the main effect of infection after accounting for treatment and the main effect of treatment after accounting for infection. The infection coefficient represented the shared transcriptional difference between EA7 and A6 under Atonik® treated conditions and between E8 and K7 under untreated conditions. Conversely, the treatment coefficient represented the shared difference between A6 and K7 under non-infected conditions and between EA7 and E8 under infected conditions. DESeq2 size factors were estimated using the median of ratios method. Gene wise dispersion estimates were calculated, a mean dispersion relationship was fitted, and the final dispersion estimates were used in negative binomial generalized linear models. The infection contrast compared infected samples EA7 and E8 with non-infected controls A6 and K7. The treatment contrast compared the Atonik® treated samples A6 and EA7 with the untreated samples K7 and E8. Log₂ fold-change estimates were stabilized using the heavy-tailed prior shrinkage method implemented in apeglm ^60^ through the DESeq2 lfcShrink() function. Multiple-testing correction was performed using the Benjamini -Hochberg false discovery rate procedure ^61^. Genes were considered differentially expressed when they met both an adjusted p-value threshold of 0.05 and an absolute shrunken log₂ fold change of at least 1. For exploratory visualization, filtered count data were subjected to variance-stabilizing transformation using vst() with blind = FALSE. PCA was performed on the variance-stabilized expression matrix using plotPCA(), with infection status and treatment included as sample annotations. Pairwise distances among libraries were calculated from the transposed variance-stabilized expression matrix and visualized using the pheatmap package^62^. Heatmaps were generated separately for the 30 genes with the lowest adjusted p-values in the infection and treatment associated contrasts (Supplementary Material S5). Variance stabilized expression values were standardized independently for each gene by calculating row wise Z-scores. Genes and samples were hierarchically clustered, and sample annotations indicated infection and treatment status. MA plots were generated from the shrunken log₂ fold-change estimates. Normalized gene counts were obtained using the DESeq2 counts() function with normalized = TRUE and exported for descriptive examination of expression patterns in the individual A6, K7, EA7, and E8 libraries. Normalized counts were used for visualization and sample-level inspection, whereas differential expression testing was based on the original integer count matrix. Gene annotations were extracted from Vitis_vinifera_GCF_030704535.1.gtf using the rtracklayer package ^63^. Records annotated as genes were selected, and the gene identifier, gene symbol, functional description, gene biotype, and NCBI database cross-reference were retained where available. The resulting annotation table was matched to the DESeq2 results using the gene_id attribute. Annotated result tables were generated separately for infection-associated genes, treatment-associated genes, and genes shared between the two contrasts. An additional categorical variable was assigned to indicate the experimental group in which each transcript showed higher expression.

Genes potentially associated with reactive oxygen species metabolism were identified by keyword searches of the gene symbols and functional descriptions. The search terms included “superoxide dismutase”, “SOD”, “peroxidase”, “catalase”, “glutathione”, “ascorbate”, “peroxiredoxin”, and “thioredoxin”. Raw RNA-seq reads have been deposited under BioProject no. PRJNA1479983 (SRA: SRR39850238, SRR39850237, SRR39850236, SRR39850235).

## Declarations

### Ethics approval and consent to participate

Not applicable.

### Consent for publication

Not applicable.

### Data availability

*In planta* experiment used *Eutypa lata* isolate E2 (MEND-F-0403), described in this study, GenBank Acc. Nos. PZ564299 (beta-tubulin gene, *Tub2*) and PZ578101 (RNA polymerase II subunit 2 (RPB2) gene). Raw RNA-seq reads have been deposited in GenBank under BioProject no. PRJNA1479983 (SRA: SRR39850238, SRR39850237, SRR39850236, SRR39850235). Processed transcriptomics result tables supporting the findings are provided in the Supplementary Material.

### Authors’ contributions

J.Č. performed the experimental work, analysed the data, prepared figures and tables, and drafted the manuscript. K.Š. contributed to the in vitro experiments, methodological design, data interpretation, and manuscript revision. P.P. performed RNA extraction, reverse transcription, qPCR analysis, and related molecular laboratory work. L.H. and D.H. contributed to plant cultivation, in planta experimental work, sample processing, and lesion measurements. M.Š. contributed to fungal isolate preparation, maintenance of fungal cultures, and methodological support. P.B. and O.D. provided the tested Atonik® formulation, contributed technical information on its application, and reviewed the manuscript. A.E. conceived and supervised the study, coordinated the project, contributed to data interpretation, funding acquisition, and manuscript revision. All authors read and approved the final manuscript.

